# Energetic cross-talk of filter gate and lower helices drives polymodal regulation and disease in TREK K2P channels

**DOI:** 10.64898/2026.02.13.705734

**Authors:** Berke Türkaydin, Valerio Rizzi, Chaimae Benkerdagh, Simon Ghysbrecht, Simone Aureli, Thomas Baukrowitz, Marcus Schewe, Francesco Luigi Gervasio, Han Sun

## Abstract

The TREK subfamily of two-pore domain potassium (K_2P_) channels are essential regulators of membrane excitability, and their activity is modulated by a wide range of physiological stimuli, including phosphorylation and membrane stretch. Single-site mutations in this subfamily have been identified in patients with FHEIG symptom (facial dysmorphism, hypertrichosis, epilepsy, intellectual disability/developmental delay, and gingival overgrowth), where they cause pathological channel hyperactivation. Using OneOPES framework, which unifies multiple enhanced-sampling molecular dynamics strategies, we provide a detailed energetic characterization of the TREK-2 conformational landscape. Our simulations uncover a unifying mechanism in which coupling between the lower transmembrane helices, the proximal C-terminal domain, and selectivity filter stability governs channel gating under different physiological stimuli, as well as disease-mimicking conditions. The predicted conformational effects of an FHEIG syndrome-associated mutation were further validated by electrophysiological measurements using a conformation-sensitive TREK-2 inhibitor. Together, these results establish an energetic framework for TREK-2 regulation and dysfunction and provide a foundation for structure-based drug discovery targeting the K_2P_ TREK channel family.

## Introduction

The TREK subfamily of two-pore domain potassium (K_2P_) channels, comprising TREK-1, TREK-2, and TRAAK, is a key regulator of neuronal excitability by mediating background leak currents that stabilize the resting membrane potential below the threshold for action potential firing^1–3^. These channels respond to a wide spectrum of stimuli including mechanical stretch, temperature, lipids, pH, and phosphorylation, demonstrating their ability to sense and integrate multiple physiological signals to modulate their activity^2,4–9^. Mechanical stretch acts as a major activating stimulus, linking TREK channel activity to changes in membrane tension that occur during neuronal firing, osmotic stress, and mechanical deformation^6,10^. Under increased membrane tension, TREK channels showed increased channel activity, leading to membrane hyperpolarization and reduced neuronal excitability. In contrast, phosphorylation of the PKC-sensitive serine residue within the proximal C-terminus (pCt) domain provides a chemical pathway for inhibition, shifting the channel toward the less-conductive down-state conformation and suppressing background leak currents^11–17^.

Based on this regulatory profile, TREK channels are implicated in processes such as neuroprotection, anesthesia, pain perception, and depression, and are recognized as promising pharmacological targets^12,18–20^. Recent genetic and electrophysiological studies have further highlighted that disruption of this regulatory balance can lead to pathological channel behavior. Gain-of-function (GoF) mutations within the TRAAK (*KCNK4*) channel have been associated with severe neurodevelopmental disorders marked by neuronal hyperexcitability^21^. Two distinct de novo missense variants (A244P and A172E) located near the M4 and M3 helices were identified in patients with FHEIG syndrome, a condition characterized by facial dysmorphism, hypertrichosis, epilepsy, intellectual disability, and gingival overgrowth. Functional analyses revealed that these mutations greatly enhance basal potassium currents and impair normal regulatory responses to mechanical or chemical stimuli, producing constitutively active channels^22^. These findings demonstrate that even single-residue mutations can disrupt gating control of TREK K_2P_ channels, leading to persistent activation and disease phenotypes.

Based on X-ray crystallographic studies, TREK and TRAAK channels adopt two major conformational states, referred to as the up- and down-states (Fig. 1a)^23–26^. These states are primarily distinguished by the orientation of their transmembrane helix M4 relative to the membrane and by the positioning of the cytoplasmic C-terminus (Ct) domain. In particular, the pCt, which lies directly adjacent to the M4 helix, has been identified as a key regulatory domain that mediates the integration of multiple intracellular signals. Notably, both the up- and down-states exhibit the same selectivity filter (SF) conformations in the crystal structures. Based on the electron density observed beneath the SF, made accessible through fenestration opening in the down-state, a lipid-based gating mechanism has been proposed to explain the functional differences between the two conformations. In parallel, electrophysiological measurements and computational simulations, including our own studies, have suggested that the SF functions as the principle gate in K_2P_ channels^7,27,28^. Furthermore, our recent study combining large-scale molecular dynamics (MD) simulations with extensive electrophysiological measurements suggested an allosteric mechanism by which the dynamics of M4 helix and pCt regulates the SF gate^17^. In down-state TREK-2 simulations, we observed pronounced backbone rearrangements near the S1 ion-binding site of the SF, which impaired ion conduction. However, this mechanism involves slow, large-scale conformational transitions within the pCt-M4 region that are difficult to capture using unbiased MD due to limited accessible timescales and the high dimensionality of the conformational landscape (Fig. 1b).

**Fig. 1.**
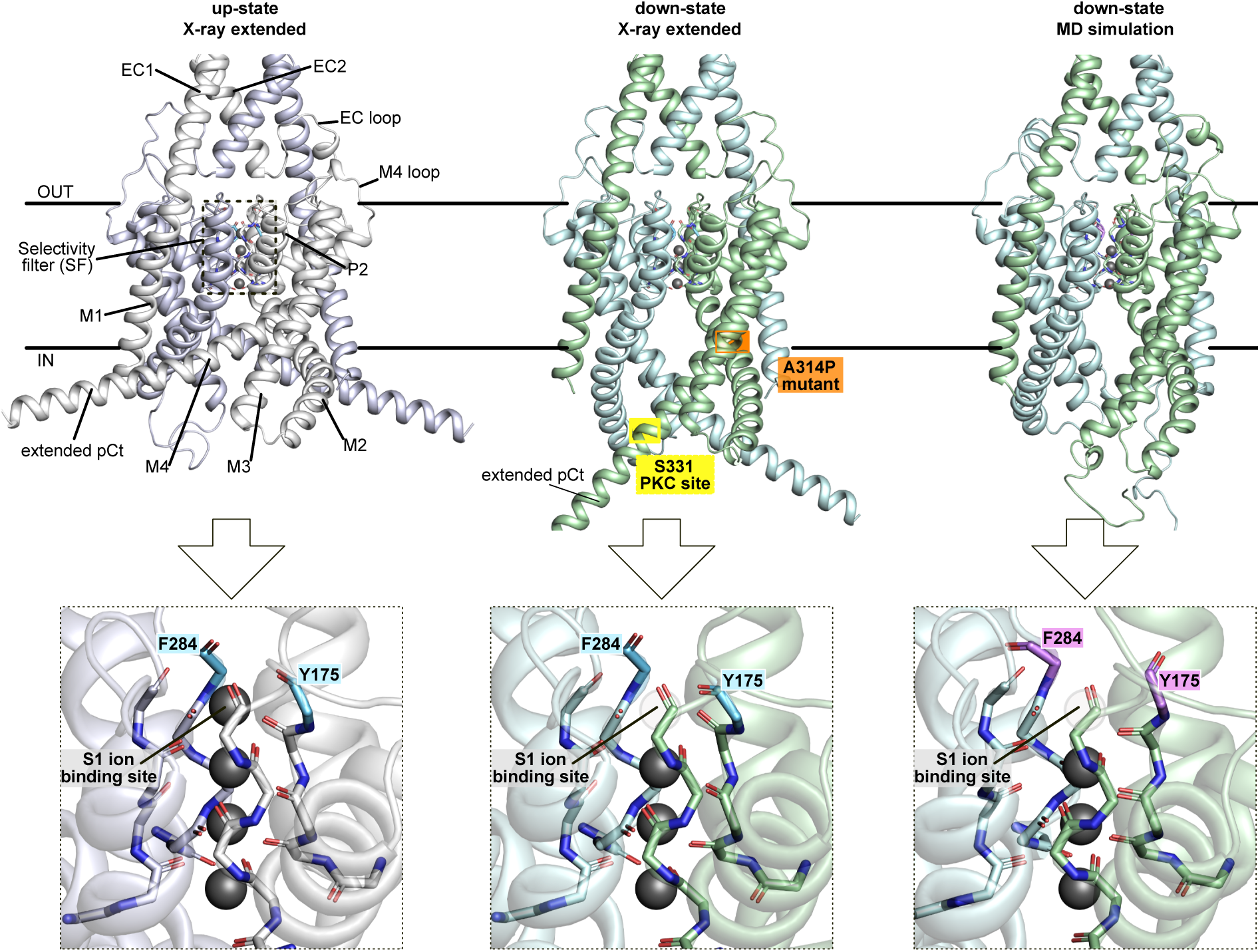
Conformational states and dynamics of TREK-2 K_2P_ channels. **(a)** Crystal structures of TREK-2 in the up-state (PDB ID: 4BW5) and down-state (PDB ID: 4XDJ)^23^ along with one representative final snapshot of TREK-2 from previous MD simulations^17^. The phosphorylation site at residue S331 in the pCt and disease mutant site are indicated with yellow and orange, respectively, as used in subsequent MD simulations. Key structural rearrangements are highlighted in cyan and purple SF residues F284 and Y175.

Over the past several decades, a number of enhanced-sampling approaches have been developed to accelerate sampling of rare conformational transitions in biomolecular systems^29–36^. However, their efficiency depends critically on how well a small number of collective variables (CVs) capture the relevant slow degrees of freedom. For large and flexible systems such as TREK channels, whose gating involves subtle long-range couplings between the cytosolic and transmembrane domains, defining an optimal low-dimensional set of CVs is particularly challenging. Standard metadynamics and similar enhanced sampling simulation methods often struggle to achieve proper convergence^30,37^. Indeed, our previous attempt using well-tempered metadynamics failed to yield a converged free energy landscape associated to the aforementioned allosteric coupling mechanism in TREK channels, leaving the energetic description of this process unresolved.

To overcome these challenges, we employed in the current study OneOPES, a recently developed extension of the On-the-fly Probability Enhanced Sampling (OPES) framework that combines adaptive biasing with replica-exchange sampling to efficiently explore high-dimensional free-energy landscapes^38^. In OneOPES, a primary replica applies an OPES Explore bias along a main collective variable (CV), targeting accurate convergence for equilibrium properties. Multiple auxiliary replicas running in parallel apply weaker, more exploratory biases along additional CVs that might involve slow motion not sufficiently accounted for by the main CV. Some replicas further incorporate an OPES MultiThermal component, which effectively heats and cools the system, lowering free-energy barriers in a manner analogous to parallel-tempering. This approach accelerates transitions along degrees of freedom not explicitly included in the CVs, mitigating the effects of suboptimal CV selection. Exchanges between replicas enable efficient sampling of rare conformations, while rigorous reweighting ensures accurate reconstruction of unbiased free-energy surfaces. These features enabled OneOPES to converge the free energy landscapes associated with protein folding and conformational changes in large and complex systems such as GPCRs and ion channels^39–41^. This makes it an ideal choice for TREK K_2P_ channels, where activation and gating are governed by allosteric network between spanning distant domains.

In this study, we applied extensive OneOPES simulations to TREK-2 to elucidate how phosphorylation, membrane stretch, and pathogenic mutations reshape the energetic landscape underlying channel gating. By combining path-based and auxiliary CVs capturing both M4-pCt and SF dynamics, we establish a robust energetic coupling between the regulatory domain and the principle gate of the channel. Our predictions of how GoF mutations in TREK-2 homologue to the pathogenic FEIGH syndrome mutations in TRAAK alter the energetic landscape were further validated by state-dependent norfluoxetine (NFx) inhibition experiments via inside-out patch-clamp measurements. Together, these findings provide a quantitative energetic framework for understanding how diverse stimuli and disease-associated mutations converge on a shared energetic coupling mechanism to modulate TREK channel activity.

## Results

### The global conformational transition of TREK-2 is driven by the coordinated motion of M4-pCt

We first selected three systems for investigations and comparisons: apo TREK-2, phosphorylated TREK-2, and a system subjected to mechanical stretch applied to the lipid bilayer. Following our previous simulations, we incorporated a 19-residue pCt into the X-ray structure^17^. For each system, we performed three independent 1 μs OneOPES simulation runs, each consisting of eight replicas (Supplementary Table 1). To characterize the overall conformational behavior of these systems, we analyzed the one-dimensional free-energy surfaces (1D FES) obtained from the OneOPES simulations, projected onto the main path CV. This reaction coordinate is able to capture the large-scale structural transition between the crystallographically resolved up- and down-state conformations of the channel (for a detailed description of the CVs, see Methods section) (Fig. 2a).

**Fig. 2.**
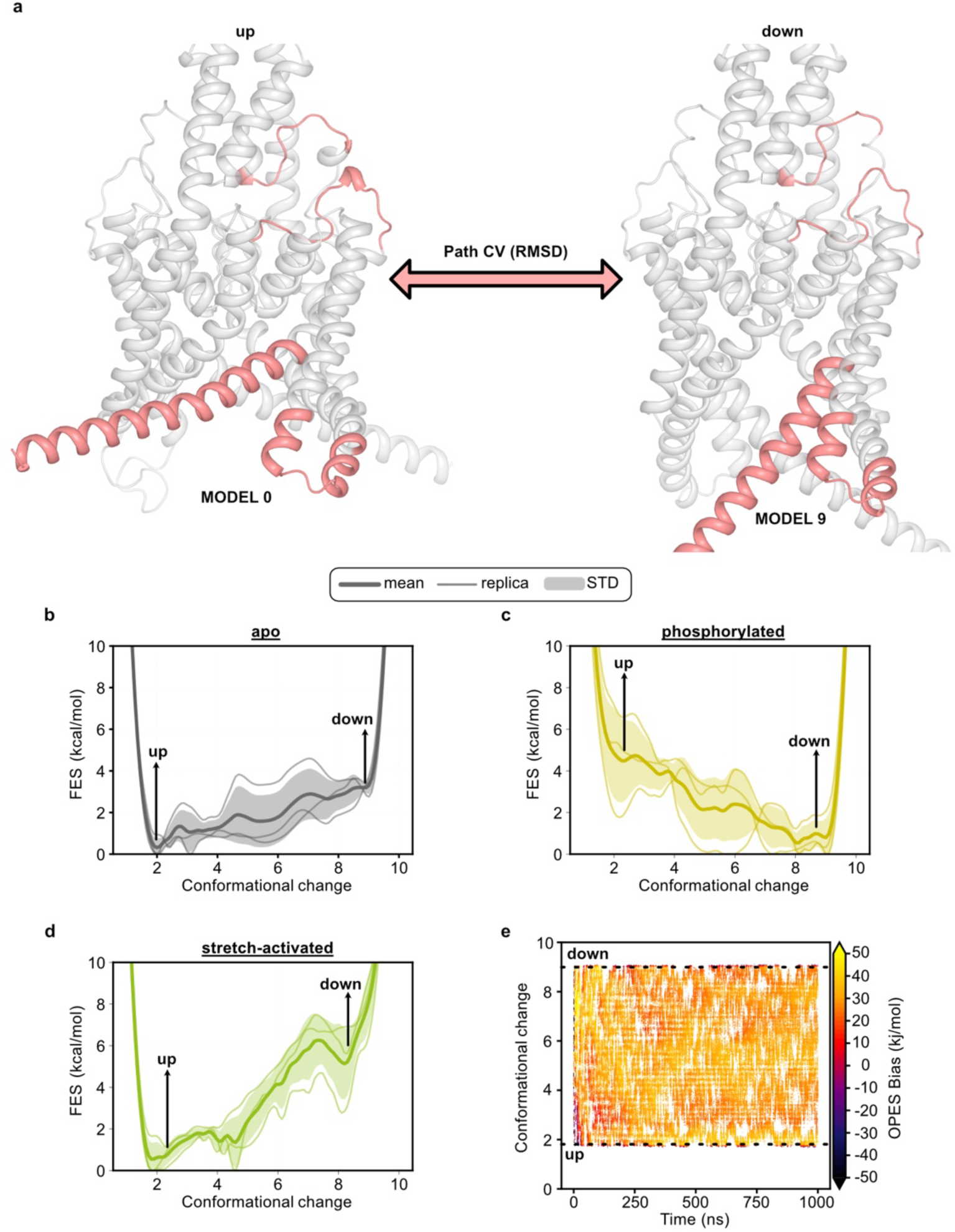
Path CV facilitates complete transition between two metastable states of TREK-2. **(a)** Representation of the preparation of path CV used in OneOPES simulations. Up-state crystal structure (PDB ID: 4BW5)^23^ was used as MODEL 0 and down-state crystal structure (PDB ID: 4XDJ)^23^ was used as MODEL 9. Highly flexible regions that are affecting the RMSD changes were colored with salmon color which were selected based on the crystal structures and observations based on our previous unbiased simulations^17^. **(b)** 1D free energy surface (FES) projected on to the path CV for apo, **(c)** phosphorylated and **(d)** membrane-stretch simulations. The thick yellow lines represent ensemble averages from three independent simulations, and the shaded area illustrates the standard deviation, demonstrating reproducibility and sampling variability. **(e)** Sampling of the path CV as a function of time in replica 0, with data points colored according to the accumulated OPES bias potential in an apo TREK-2 OneOPES simulation.

The averaged 1D FES of the apo TREK-2 system displayed a relatively flat profile with the up-state stabilized by approximately 3 kcal/mol relative to the down state (Fig. 2b). This implies that the up-state is the dominant conformation in the absence of external stimuli, while the down-state remains thermally accessible. In the phosphorylated TREK-2 system, the FES shifted to a thermodynamically favored down-state (Fig. 2c), with an energy difference of about 4.7 kcal/mol. This is in line with the experimentally observed inhibitory effect of phosphorylation^17^. By contrast, mechanical stretch drastically altered the energy profile (Fig. 2d). In this system, the 1D FES collapsed into a single dominant minimum corresponding to the up-state, while the down-state is strongly disfavored. The free-energy difference between the two conformations exceeded 6 kcal/mol, suggesting strong thermodynamic bias towards the up-state conformation under mechanical activation. These results are fully consistent with previous experimental and computational studies that identified mechanical stretch as a key regulator of TREK channel activation^10,42^.

Convergence analysis of the OneOPES bias potential and the time evolution of the path CV further confirmed the robustness of sampling across all replicas (Fig. 2e and Supplementary Fig. 1). Each simulation achieved stable bias deposition and consistent free-energy estimates within the first several hundred nanoseconds, indicating efficient exploration of the relevant conformational space. These results indicate that OneOPES accurately captures the essential gating motions of TREK-2 and provides a quantitatively converged description of its conformational energetics across different regulatory conditions.

To examine in greater details how the global up-down transitions are coupled to the structural rearrangements of the M4 helices, we next analyzed the two-dimensional free-energy surfaces (2D FES) spanned by the path CV and the two auxiliary CVs describing relative M4–M2 displacements (Fig. 3a). In a previous study, the pCt-M4 region was shown to undergo a pronounced vertical and lateral movement that redefines its interactions with the adjacent M2 helices^17^. Accordingly, the auxiliary CVs were specifically designed to quantify the relative displacement of the M4 helices with respect to the M2 helices. One auxiliary CV (AdA) tracks the distance between the M4 and M2 helix within the same subunit, which decreases as the channel moves towards the down-state. The second CV (BdA) monitors the distance between the M4 and M2 helix of the opposite subunit, which instead decreases towards the up-state.

**Fig. 3.**
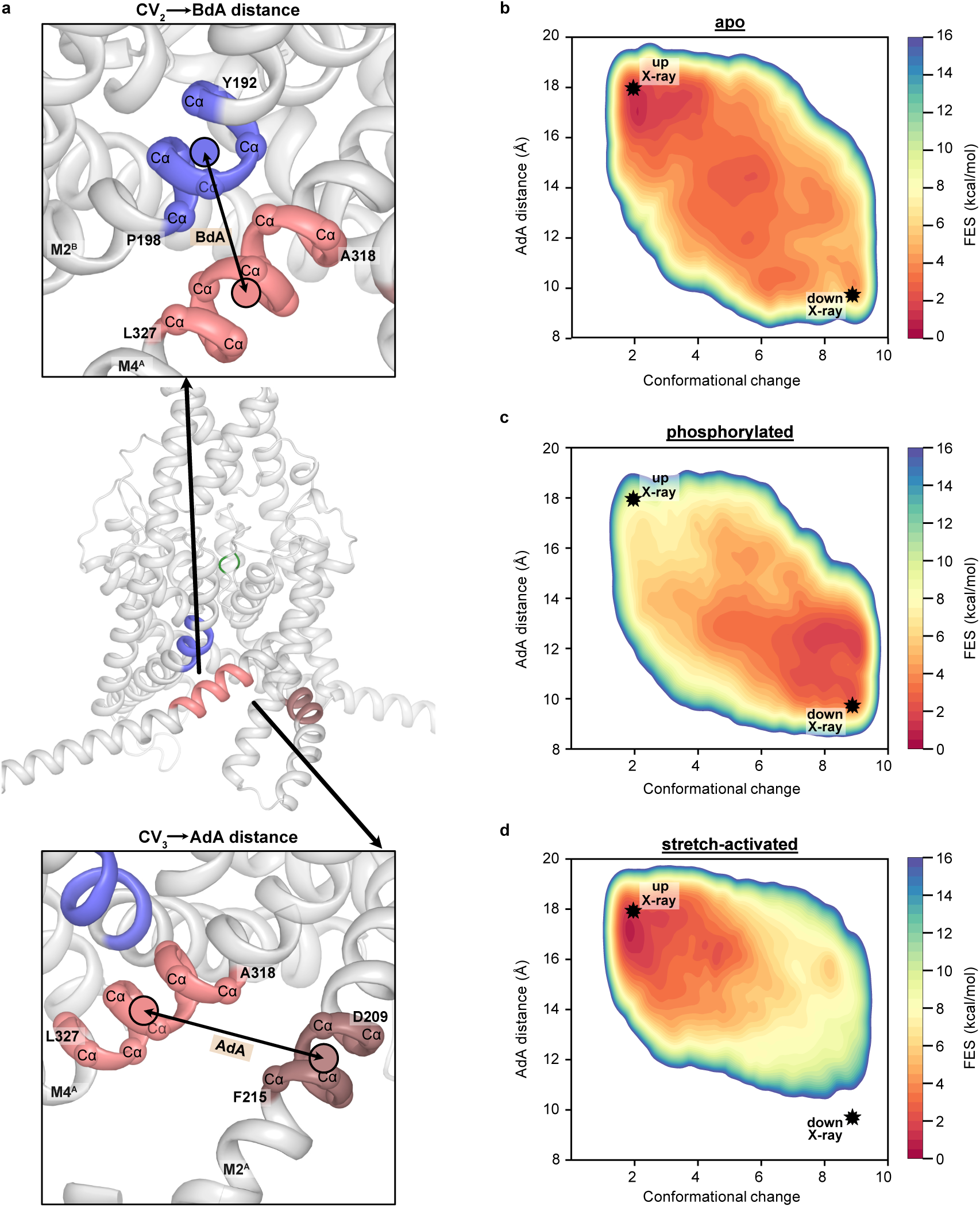
Structural rearrangements of the pCt provides energetic link to overall conformational dynamics. **(a)** Representation of the two auxiliary CVs describing the pCt rearrangement in up- and down-states. The center of mass (CoM) distances of Cα atoms between selected residues are used to measure distance between M4-M2 helices for M4 and M2 located in the same chain A (AdA distance) and M4 and M2 of opposite chains (BdA distance). **(b)** Two dimensional free energy surfaces projected along the path CV and the auxiliary CV describing M4-M2 distance for apo, **(c)** phosphorylated and **(d)** membrane-stretch simulations. The average of three independent replica for each system were calculated for two dimensional free energy surfaces. Black stars indicate the positions of the crystallographic TREK-2 up- and down-state structures mapped onto these coordinates, based on their measured path CV values and AdA distances.

Across all systems, the 2D FES (Fig. 3) underscore again a strong correlation between the M4-M2 displacement and the path CV: low path CV values (up-state) coincide with high AdA and low BdA, whereas high path-CV values (down-state) correspond to the opposite configuration (Fig. 3b and Supplementary Fig. 2). This tight coupling indicates that the auxiliary CVs faithfully track the structural rearrangements that accompany the global gating motion. Consistent with the 1D FES, the apo and the membrane-stretched systems show a stabilized up-state, while the phosphorylated system shows a stabilized down-state. In the apo system, the 2D FES is relatively shallow and broadly dispersed across both CV axes, reflecting a more permissive and dynamically heterogenous motion of the M4 helix under equilibrium conditions. By contrast, phosphorylated and membrane-stretched systems exhibit markedly more localized free energy basins, with population concentrated around a single, well-defined minimum. These compact landscapes indicate a substantial restriction of M4-M2 rearrangements: phosphorylation constrains the helix toward the down-state geometry (with M4 closely aligned to M2 of the same subunit), whereas membrane stretch confines the system to the up-state (with M4 drawn towards M2 of the opposing subunit). Together, these results revealed that the main path CV and M4-related auxiliary CVs consistently captured the global structural rearrangements of TREK-2 across different regulatory conditions and demonstrated that phosphorylation and membrane stretch act as opposing modulators of the same energetic pathway.

### Allosteric communication between the pCt and SF defines the energetic basis of TREK-2 gating

After establishing the energetic correspondence between the path CV and dynamics of the M4 helices, we next examined whether these conformational transitions at the M4-pCt influence the local gating behavior of the SF. As discussed before, the SF represents the narrowest region of the pore and has been proposed to serve as the primary gate controlling ion conduction in TREK channels. Our previous work has shown that subtle dihedral rotations within the SF backbone at the S1 K^+^ binding site (Fig. 4a) can drive TREK channels into a non-conductive conformation^17^. To capture this transition in energetic terms, we projected the OneOPES free-energy surfaces along the path CV and two auxiliary CVs corresponding to the ψ dihedral angles of residues F284 and Y175 within the SF (Fig. 4a). These coordinates describe the structural rearrangements occurring at the S1 K^+^ binding site and allow quantitative assessment of the coupling between the M4-driven conformational dynamics and SF gating.

**Fig. 4.**
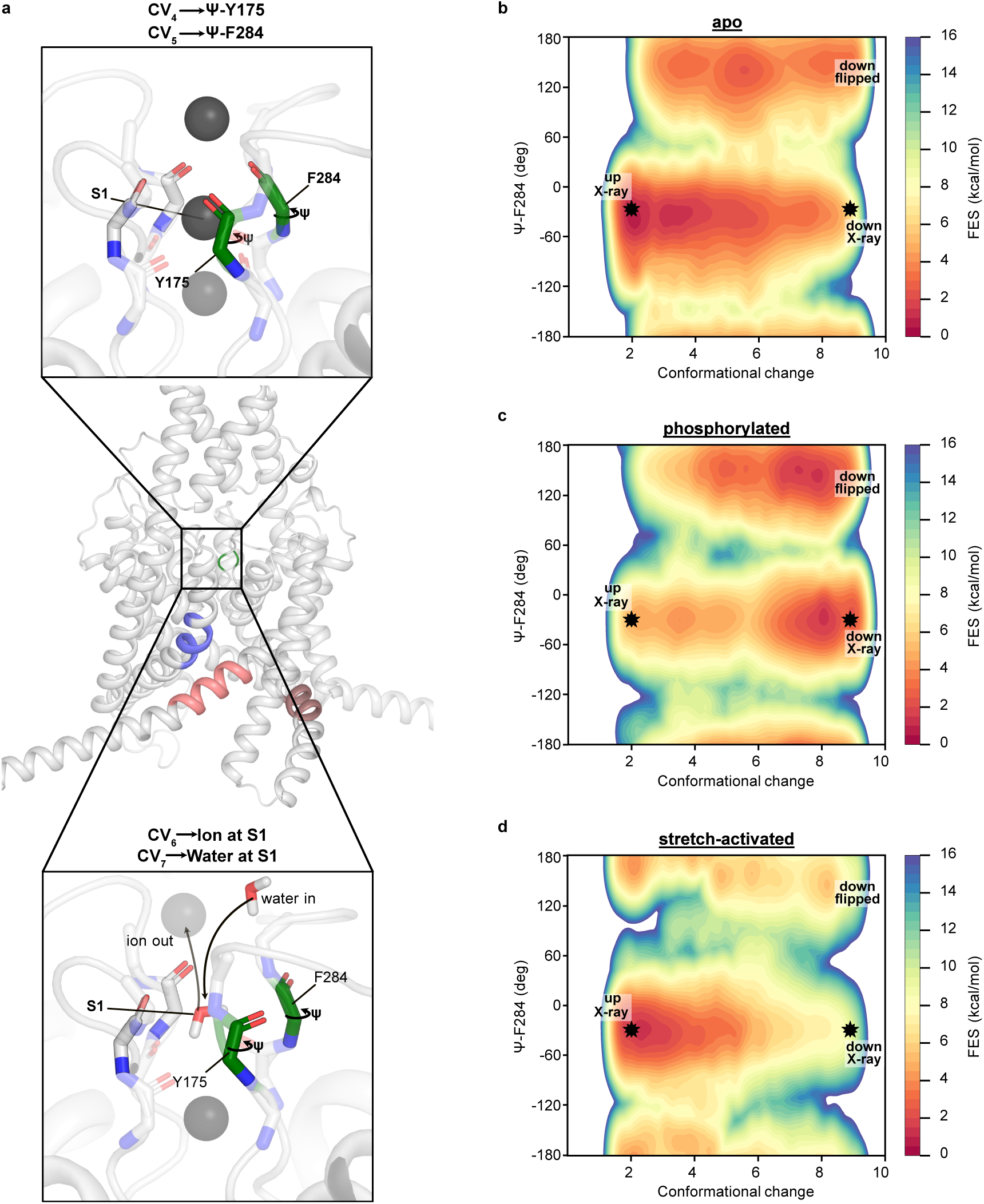
Auxiliary CVs to describe SF dynamics provide energetics of allosteric coupling between conformation of TREK-2 channel and gating. **(a)** Representation of the two auxiliary CVs describing the SF stability in conductive and non-conductive conformations. Two additional auxiliary CVs were also used to describe the ion and water occupancy at the S1 ion binding site further helping the transition of SF residues. **(b)** Two dimensional free energy surfaces projected along the path CV and the auxiliary CV describing SF stability for apo, **(c)** phosphorylated and **(d)** membrane-stretch simulations. The average of three independent replica for each system were calculated for two dimensional free energy surfaces. Black stars indicate the positions of the crystallographic TREK-2 up- and down-state structures mapped onto these coordinates, based on their measured path CV values and AdA distances.

In the apo simulations, the up-state region of the energy landscape predominantly coincided with a conductive SF configuration, where the ψ dihedral angle of F284 remained in their ion-coordinating crystallographic state (Fig. 4b). A metastable flipped-SF conformation was also observed within the up-state region, about 3 kcal/mol higher in energy. Importantly, the barrier separating canonical and flipped SF in the up-state region is large (∼10-12 kcal/mol), consistent with the absence of SF flipping in our previous unbiased simulations of the up-state conformation. As the channel moves toward the down-state, the canonical SF is destabilized, while the flipped SF remains comparatively constant and eventually becomes the thermodynamic minimum. Concurrently the barrier between the two SF conformers decreases (to ∼6 kcal/mol) such that flipping becomes considerably more accessible and conformations with a flipped SF, corresponding to the non-conductive state, become dominant. Together these results support the view that, under basal conditions, TREK-2 dynamically exchanges between conductive and non-conductive states while maintaining an overall equilibrium between the two global conformations. Here, M4-pCt motion both thermodynamically biases the SF equilibrium and modulates the local barrier to carbonyl flipping, providing a mechanistic basis for allosteric control of SF gating.

In addition to F284, we also monitored the ψ dihedral angle of Y175, located on the opposite side of the S1 site. Although Y175 represents an important structural component of the SF, its sampling behavior in our simulations deviated from that observed for F284 Supplementary Fig. 3a). We observed that both SF conformers were thermodynamically accessible within the up-state region of the free-energy landscape, a feature not sampled in our previous unbiased simulations. This difference likely arises from the limited set of auxiliary CVs used to describe Y175 dynamics. In our previous unbiased simulations, destabilization of Y175 was driven by interactions between the M4 loop and the extracellular (EC) loop; however, these collective variables were not explicitly included here, resulting in incomplete thermodynamic sampling of Y175 flipping.

Upon phosphorylation, the up-state became destabilized, and the phosphorylated channel predominantly occupied the down-state, where SF flipping was more energetically favored (Fig. 4c and Supplementary Fig. 3b). This result provides further energetic validation that the dynamics of the SF and M4 helices are allosterically coupled to one another.

Conversely, the stretch-activated system displayed the opposite energetic profile. Under mechanical stretch, the energy landscape collapsed onto a single deep minimum corresponding to the up-state with a stable SF conformation for F284 (Fig. 4d) while again both SF conformers were thermodynamically accessible within the up-state region of the free-energy landscape for Y175 (Supplementary Fig. 3c). The barrier separating the stable and backbone-flipped SF configurations exceeded 14 kcal/mol, making conformational transition highly improbable. As a result, the SF remained predominantly in the conductive SF conformation throughout the simulations, while down-state or backbone-flipped conformations were only rarely sampled. This pronounced energetic stabilization of the up-state and the stable conductive SF conformation provides a clear mechanistic explanation for how membrane stretch promotes channel activation and prevents transition to the non-conductive state.

The multidimensional free energy landscapes obtained from the OneOPES simulations capture the full energetic spectrum of TREK-2 gating, ranging from dynamic equilibrium in the apo channel, to strong inhibitory bias upon phosphorylation, and robust activation under membrane stretch. The consistent correlation between the up-down state transitions and SF stability demonstrates that the coupling between the M4-pCt and SF dynamics represents a main energetic axis through which diverse regulatory signals modulate TREK channel function.

### Disease-mimicking mutation reshapes the TREK-2 energy landscape toward a conductive state

To investigate how a disease-associated mutation reshapes the conformational landscape of TREK-2 leading to channel dysfunction, we performed and additional series of free energy simulations with OneOPES. We selected a pathogenic gain-of-function (GoF) mutation in TRAAK (A270P) identified in patients with FHEIG syndrome (Fig. 5a) and introduced the homologous mutation (A314P) into the M4 helix of TREK-2^21^. As expected, this substitution also produced a GoF phenotype in TREK-2. Currents recorded from oocytes expressing the mutant channel were increased by 9-fold compared to wild-type WT) (Fig. 5b,c). In parallel, we performed five independent OneOPES simulation runs, each compromising 8 replicates of 1 μs, to evaluate its energetic and structural effects.

**Fig. 5.**
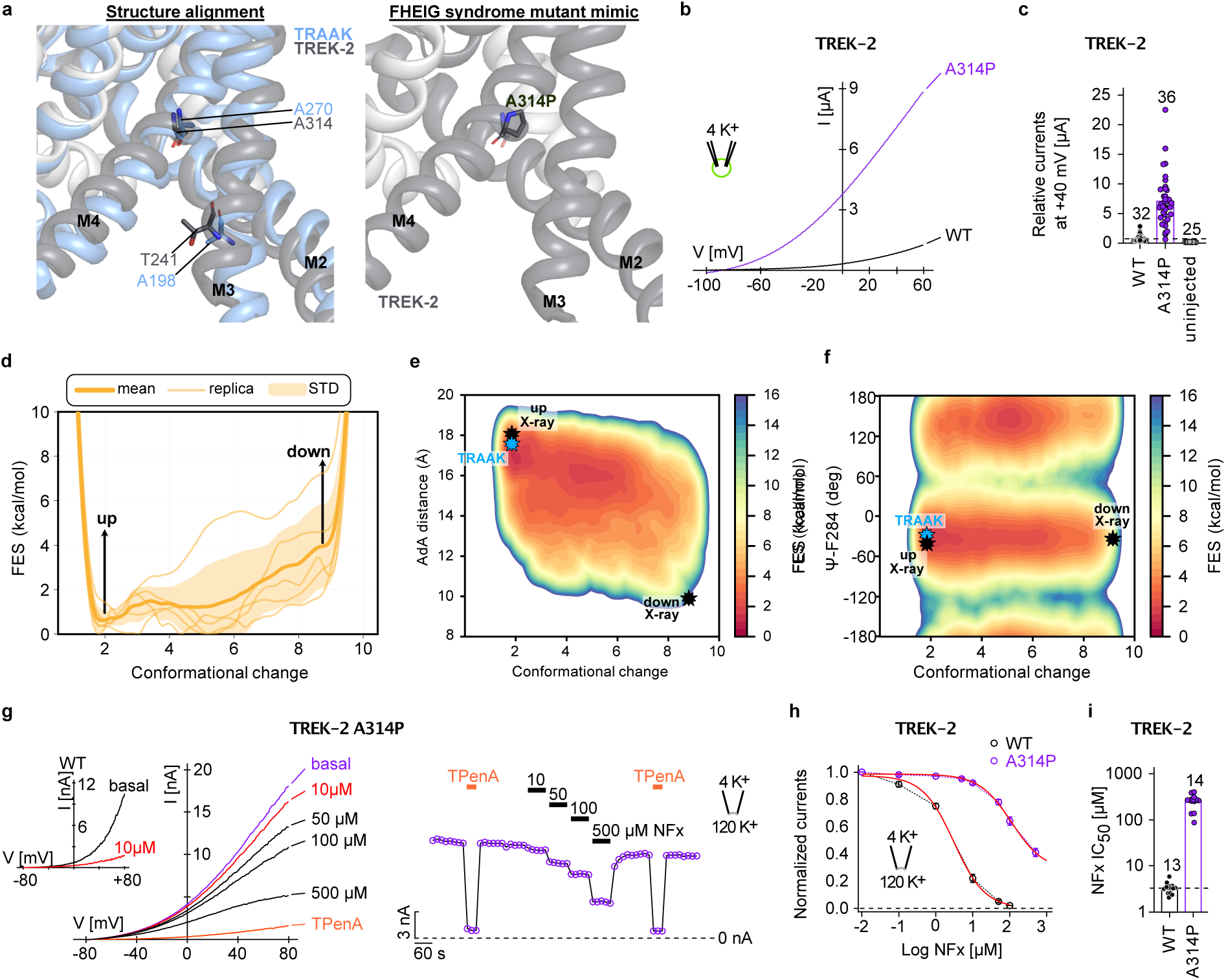
Disease-mimicking mutation stabilized TREK-2 into a conductive up-state conformation. **(a)** Alignment of two K_2P_ channels, TRAAK (PDB ID: 4WFE) and TREK-2 (PDB ID: 4BW5)^23^ with GoF mutation residues highlighted. **(b)** Example measurements and **(c)** relative currents measured from oocytes expressing TREK-2 WT, A314P mutant channels and uninjected oocytes, respectively in TEVC with extracellular low K^+^ [2 mM] at pH 7.4 and analyzed at +40 mV. Analyzed electrophysiological data shown as mean ± SEM from n (number of independent experiments) as indicated in the figure panel. **(d)** 1D free energy surface (FES) projected on to the path CV for disease-mutant (A314P) TREK-2 simulations. **(e)** Two dimensional free energy surfaces projected along the path CV and the auxiliary CV describing M4-M2 distance in the opposite subunit (BdA distance). **(f)** Two dimensional free energy surfaces projected along the path CV and the auxiliary CV describing SF stability for disease-mutant TREK-2 simulations. An X-ray structure of previously determined TRAAK A270P mutant (PDB ID: 7LJ4)^43^ was projected onto the 2D FES as blue star. Black stars indicate the positions of the crystallographic WT TREK-2 up- and down-state structures mapped onto these coordinates, based on their measured path CV values and AdA distances. **(g)** Example measurements of WT TREK-2 and A314P mutant channels in an asymmetric K^+^ gradient at pH 7.4 in a voltage range from -80 to +80 mV showing the dose-dependent inhibition with the indicated concentrations NFx (left) and the time course of block analyzed at +40 mV (right). **(h)** Dose-response curves of NFx inhibition for WT and mutant TREK-2 channels as indicated analyzed from recordings as in (g). **(i)** NFx IC_50_ values from dose-response curves as in (h) for WT and mutant TREK-2 K_2P_ channels.

Both one- and two-dimensional free-energy surfaces projected along the path CV and the auxiliary CVs describing M4 dynamics revealed a distinct energetic profile compared to the WT system (Fig. 5d-f and Supplementary Fig. 2d). In contrast to the shallow, more broadly distributed landscape of the apo channel, the mutant exhibited a tightly localized energetic basin centered on the up-state conformation. Similar to the membrane-stretched system, the 2D FES shows the M4 helix drawn toward the M2 helix of the opposing subunit (low BdA value), accompanied by markedly reduction in conformational freedom. However, relative to the membrane-stretched condition, the TREK-2 A314P simulations exhibit slightly broader fluctuations within the up-state basin, indicating increased conformational flexibility of the M4-pCt dynamics.

Energetic and structural inspection provides the mechanistic basis of this stabilization. The proline substitution introduced a kink within the M4 helix which alters the geometry of the pCt-M4 interface, forcing it to adopt a higher, laterally displaced position closer to the opposing M2 helix. As a consequence, the channel was stabilized in the up-state even in the absence of external force, indicating that the single-residue mutation effectively mimics the energetic effect of stretch activation.

To experimentally probe the down-up-state equilibrium of TREK-2, we used the state-dependent inhibitor norfluoxetine (NFx), which binds within the side-fenestrations that are accessible only in the down-state and thereby inhibits channel activity^23,44,45^. NFx was applied at increasing concentrations to the cytoplasmic side of excised membrane patches expressing WT or mutant (A314P) TREK-2. The mutation shifted the IC₅₀ for NFx inhibition from 3.4 ± 0.3 µM (WT) to 256.7 ± 23.4 µM (A314P), consistent with a pronounced shift of the conformational equilibrium toward the NFx-insensitive up-state (Fig. 5g-i). Furthermore, mapping the previously determined crystal structure of the corresponding TRAAK A270P mutant onto the 2D free-energy surface (FES) of the TREK-2 simulations demonstrates that the energetic minimum identified in the TREK-2 landscape closely aligns with the experimentally resolved TRAAK mutant structure (Figure 5)^43^.

We next examined whether this stabilization of the up-state affected the energetics of the SF. Projections of the free-energy surfaces along the Path CV and the auxiliary CV describing the ψ dihedral angle of F284 revealed that the conductive SF conformation was sampled effectively, indicating that the channel consistently remained in its conductive orientation under membrane stretch (Fig. 5e). Similar to the apo simulations, the ψ dihedral angle of Y175 showed incomplete thermodynamic sampling, likely due to the absence of CVs that capture the underlying structural drivers of this dynamics (Supplementary Fig. 3d). The free-energy barrier separating the conductive and backbone-flipped conformations exceeded ∼8 kcal mol⁻¹, making conformational changes of the SF highly improbable under these conditions.

Together, these observations demonstrated that the disease-mimicking mutation A314P stabilized the up-state and the conductive conformation of the SF, resulting in a high-conductive, gain-of-function-like phenotype. The energetic profile of the mutant closely resembled that of the stretch-activated system.

## Discussion

TREK channels are polymodal K^+^ channels regulated by a diverse range of external stimuli^2,3,6,7^, and their dysfunction has been implicated in multiple neurological disorders^13,46–48^. In this work, we establish a comprehensive energetic framework that elucidates how TREK-2 gating is modulated by phosphorylation, mechanical stretch, and a *de novo* mutation of the corresponding TRAAK channel associated with FHEIG syndrome (Fig. 6). We show that, although these regulatory inputs act at distinct structural sites − phosphorylation at the pCt, mechanogating via membrane-pCt/M4 coupling, and the disease-associated mutation within the M4 helix − they converge on and reshape the same underlying conformational landscape that governs channel gating. By employing an extensive set of CVs within the recently developed OneOPES enhanced sampling framework^38^, we explored TREK-2’s functional dynamics and obtained converged free energy landscapes that capture the complex allosteric coupling between M4-pCt motions and the SF gating. This approach bridges the gap between static structural determination and dynamic physiological regulation, revealing how distinct external stimuli and a pathogenic mutation reshape a shared energetic landscape to modulate channel activity and dysfunction.

**Fig. 6.**
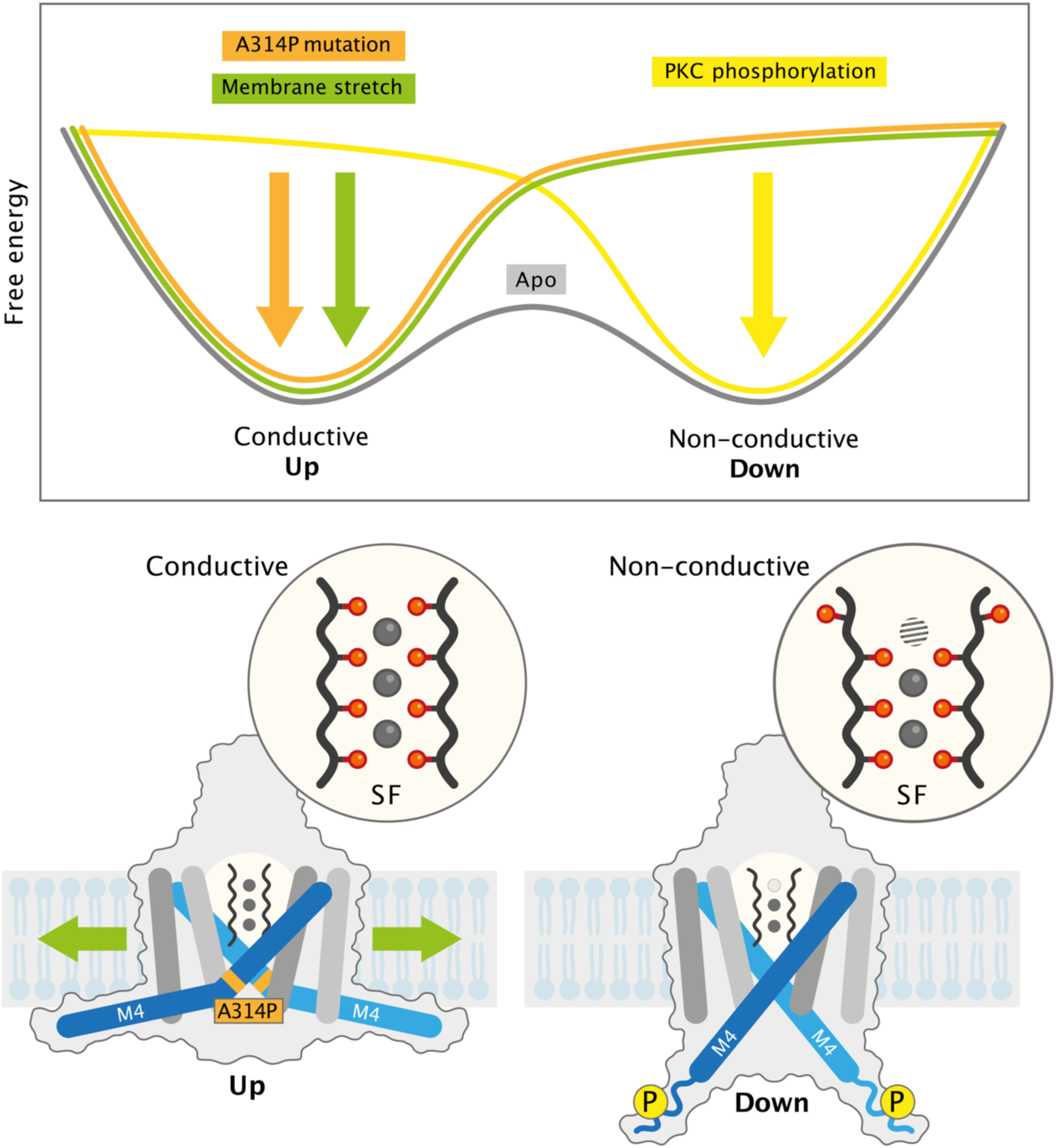
Energetic modulation of TREK-2 gating by phosphorylation, membrane stretch, and disease-associated mutation. Schematic representation of the free-energy landscapes describing TREK-2 conformational equilibria. In the apo state, the channel exhibits a bimodal energy profile with comparable up- and down-state minima. Posttranslational phosphorylation at the PKC site shifts the equilibrium toward the non-conductive down-state, whereas membrane stretch and the A314P mutation both favor stabilization of the conductive up-state. Lower panels illustrate the corresponding structural and functional outcomes: upward displacement of the M4 helix and a stable, ion-conductive SF under activating conditions, versus a downward conformation and disrupted filter in the inhibited, phosphorylated state.

The reliability of the OneOPES results rests on a multidimensional CV design that captures both global and local determinants of gating. In addition to the path CV describing the overall up-down transition, we introduced six auxiliary CVs representing distinct dynamic features of the channel as suggested by our previous unbiased MD simulations starting from up- and down-state of TREK-2 channel, respectively^17^. Two CVs quantified the relative displacement between the M4 and M2 helices within and across subunits, providing an energetic measure of the pCt–M4 motion coupled to conformational change. Two CVs tracked the dihedral angles of key SF residues (F284 and Y175) to monitor conductive to non-conductive transitions at the filter, while two additional CVs described ion and water occupancy at the S1 site, reflecting local stabilization of the gating configuration. Together, these CVs enabled OneOPES to treat both large-scale domain rearrangements and local gating events in a unified energetic landscape, which is difficult to achieve with conventional enhanced sampling methods.

The resulting free-energy surfaces showed that TREK-2 operates through two states of the lower helices corresponding to crystallographic up and down conformations. In the apo system, free-energy landscapes were shallow, allowing dynamic exchange between them and resulting in alternating conductive and non-conductive configurations of the SF. Analysis of the free-energy landscape also revealed that the up-state configuration does not correspond to a static conformation but rather encompasses a dynamic ensemble of up-state-like structures. Within these conformations, the SF remains in a stable, conductive configuration as revealed in the crystal structure. Phosphorylation at the pCt shifted the equilibrium towards the down-state and increased population in non-conductive SF configurations. This finding aligns with our previous computational and experimental observations showing that phosphorylation promotes conformational changes at the SF and inhibits ion conduction^17^. In contrast, the application of membrane stretch stabilizes the up-state, and the channel predominantly sampled conductive configurations of the SF throughout the simulations.

In this study, we obtained a good correlation between the global M4-pCt displacement and the stability of F284 at the S1 site. In contrast, the energetic description of the Y175 deviates from our previous unbiased simulations, which we attribute to the absence of additional CVs capturing conformational changes in the M4 loop and extracellular loop that contribute to Y175 rearrangements. Furthermore, unlike our earlier unbiased simulations performed using the Computational Electrophysiology framework^17^, the current setup does not include transmembrane voltage, which may further differ in the sampling of Y175 conformational transitions. Despite these limitations, incorporating Y175 as a CV proved critical for improving the overall description of SF flexibility and for facilitating the proper sampling of F284 rearrangements, analogous to the use of auxiliary CVs describing ion and water occupancy at the S1 site.

Atomistic mechanism of mechano-gating in TREK channels has been a central topic in previous computational studies employing both all-atom and coarse-grained simulations^10,42,49^. Coarse-grained simulations have primarily focused on achieving sufficient sampling of the up-to-down transition in the M4 helix of TRAAK channel by applying surface tension, in line with the “force from lipid” principle of mechanosensitivity^49^. In parallel, several all-atom MD studies on TREK-2 reported SF conformational rearrangements and associated changes in ion occupancy within the filter. However these unbiased MD simulations, limited by their accessible timescales, did not establish an energetic link between the SF dynamics and conformational transitions in the M4 helix. We noted that the specific SF rearrangements reported in these studies differ. While the MD study by J. T. Brennecke *et. al* proposed conformational changes at the S3 site^42^, P. Aryal and we independently observed up- and down-state-dependent rearrangements at the S0 and S1 sites^10,17^. We do not view these results as contradictory, as our previous MD study on non-canonical voltage gating suggested that conformational changes at the S1 and S3 sites in TREK channels are coupled to each other^50^.

Notably, TREK channels are also modulated by temperature, and recent studies have shown that thermosensitivity is regulated through interactions between the C-terminus and microtubule adaptor proteins^51^. Based on the present findings, we propose that this interaction may modulate the dynamics and orientation of the Ct and M4 helix, thereby coupling to the SF gate and fine-tuning channel activity.

A disease-mimicking mutation (A314P) in TREK-2 offered a distinct perspective on the underlying energetic landscape. Substitution of the conserved A314 residue with proline within the M4 helix, analogous to a gain-of-function mutation previously reported in TRAAK channels^21^, resulted in a pronounced energetic basin corresponding to the up-state, while the down-state was energetically inaccessible across all replicas. Structural inspection revealed that the proline substitution introduced a subtle kink in M4 that increased local flexibility and relieved steric strain at the pCt-M4 interface. This conformational relaxation favored an expanded geometry of the channel, closely resembling the mechanically stretched condition. Notably, mapping the previously determined X-ray structure of the corresponding TRAAK mutant onto the 2D FES demonstrated that the energetic minimum from the TREK-2 A314P simulations aligns closely with the experimental TRAAK mutant structure. This finding further suggests that our computational framework could be extended to other *de novo* TRAAK mutations associated with similar FHEIG phenotypes. These additional mutations also cluster within the intracellular M2-M3-M4 interaction network, which is expected to influence the up-to-down transition^43^.

Together, these findings emphasize that TREK-2 gating is governed by a finely tuned energetic coupling network connecting the pCt, M4 helix, and SF. Phosphorylation, membrane stretch, and mutation modulate different elements of this network, shifting the balance between M4 up- and down-state populations and thereby directly controlling ion conduction at the SF gate. Although lipid block enabled by membrane fenestration has been proposed to influence the TREK channel conductance^23^, our study provides an energetic evidence for the allosteric coupling between distal domains and suggests a mechanism by which the functional dynamics of the M4 helix can modulate the conductance even in the absence of lipids. The ability of OneOPES to accurately reproduce these shifts demonstrates its power for describing high-dimensional, slow conformational processes in complex membrane proteins. Incorporating multiple auxiliary CVs that captured both global conformational changes and local gating rearrangements proved essential for achieving convergence and mechanistic interpretability.

In summary, our simulations establish a coherent energetic model that explains how chemical, mechanical, and mutational perturbations modulate TREK-2 function through shared structural pathways. The interplay among pCt dynamics and their membrane interactions, M4 conformational transitions, and SF stability defines the channel’s gating landscape. The OneOPES framework enables direct connection of these slow, collective motions to allosteric regulation. These insights provide a foundation for the rational design of pharmacological strategies aimed at precisely modulating these processes. It also pays the way for future integrative studies combining simulations, mutagenesis, and electrophysiology to unravel the molecular logic underlying polymodal gating in other ion channels.

## Methods

### MD-based system preparation

The apo-, phosphorylated, stretch and disease-mutant structures of TREK-2 were generated starting from up- (PDB ID: 4BW5)^23^ and down-state (PDB ID: 4XDJ)^23^ conformations, respectively. The missing pCt region in each subunit was manually extended using the PyMOL builder tool^52^. To maintain the α-helical conformation, the “helix” setting was applied during the sequential addition of residues. A total of 19 residues (333 - 351: TKEEVGEIKAHAAEWKANV), corresponding to the Ct extension from the TREK-2 construct^53^, were appended. N- and C-terminal caps were added using N-methylamide and acetyl groups, respectively, also via the PyMOL builder. Additionally, missing loop segments (residues 149–154 and 229–235) were modeled using SWISS-MODEL^54^.

We performed MD simulations with AMBER99sb^55^ force field. The insertion of the TREK-2 channel into a POPC was performed with the GROMACS internal embedding function. The concentration of KCl was 600 mM in simulations, in consistent with our previous unbiased MD simulations under transmembrane voltages^17^, where higher ionic concentration compared to physiological condition was applied to accelerate ion conduction rate. Improved ion parameters^56^ and lipid parameters^57^ were employed in the simulations. The SPC/E water model^58^ was used in simulations.

All MD simulations were performed with the GROMACS software (2022.5)^59^ patched with PLUMED 2.8.5^60^. Short-ranged electrostatics and van der Waals interactions were truncated at 1.0 nm. Long-range electrostatics were calculated with the Particle-Mesh Ewald summation^61^. Temperature and pressure coupling were treated with the V-rescale^62^ scheme and the Parrinello-Rahman barostate^63^, respectively. The temperature was set to 300 K and the pressure to 1 bar. Fluctuations of the periodic cell were only allowed in the z-direction, normal to the membrane surface, keeping the density of the membrane unchanged. Verlet cut-off scheme was used for neighbor searching and pair interactions^64^. All bonds were constrained with the Linear Constraint Solver (LINCS) algorithm^65^.

After the channel was embedded into POPC with ions, the system was energy minimized and equilibrated. The energy minimization was done with GROMACS “steepest descent” algorithm with a maximum energy value of 100 kJ mol^-1^ to make sure to minimize the energy of the system as much as possible. After the minimization, the system was equilibrated for 10 ns with a position restraint by a force constant of 1000 kJ mol^-1^ nm^-2^ in x, y, and z-direction on the backbone atoms of the protein followed by 20 ns without position restraints on the backbone atoms of the protein (Supplementary Fig. 4). During the complete equilibration, the isothermal–isobaric (NPT) ensemble was used.

The simulations were initiated with a four-ion configuration (S1 - S4) in the SF for all setups, consistent with the ion configuration in the X-ray structure of the TREK channel in the up-state (PDB ID: 4BW5)^23^.

For the in-silico preparation of the phosphorylated serine residue in the TREK-2 in up-state structure, the hydrogen of the hydroxyl group was removed, and a phosphate group was added to the oxygen using the builder option of PyMOL. The hydrogen atom in the phosphate group was removed to maintain the phosphoserine at a -2 charge. Additional force field parameters for phosphorylated serine were adopted from previous studies^66^.

To investigate the stretch-induced changes in TREK-2, the bilayer plane (xy-plane) pressure was applied (lateral tension) at -50 bar, whereas the pressure in bilayer normal (z) direction kept at +1 bar and the temperature at 300 K. To measure the lipid bilayer thickness and the stability of the system, the volume of the POPC bilayer was calculated during the simulation time and the center-of-mass (COM) distance of head groups of the bilayer was monitored. Same protocol was followed for the rest as the apo system.

For the in-silico preparation of the disease-related mutant in the TREK-2 in up-state structure, the mutation of the Ala314 to Pro was performed using the mutagenesis tool of PyMOL^52^. Same protocol was followed for the rest of the preparation of the simulation system similar to the apo system.

### OneOPES simulations

We adopted a CV-based enhanced sampling method to investigate the energetic conformational landscape of apo-, phosphorylated, membrane stretch, and disease-mutant TREK-2 structures. We employed “On-the-fly probability enhanced sampling” (OPES) algorithm in its “Explore” flavor^67^. To perform complete transitions between up- and down-state structures of TREK-2, we employed a combination of CVs. The main CV was a path CV (pCV)^68^ describing the transition of Cα atoms between the crystallographic up- and down-state structures of TREK-2. Intermediate structures for the path were generated using the PyMOL morph function, yielding 10 protein conformations (MODELS) equally spaced in RMSD between the up- (MODEL=0) and down-state (MODEL=9). Flexible regions were excluded from the alignment procedure (Fig. 1a) and used for conformational path calculation. While this CV is sufficient to drive the overall motion of the channel, we used additional auxiliary CVs to govern sampling of slow, localized gating motions, as well as ions and water occupancies in the selectivity filter:

- **CV_2_**: the distance between M2 and M4 of the same subunit. To describe the CV we selected center of mass of all Cα atoms of residues from D144 to F150 for M2 as first point and center of mass of all Cα atoms of residues from A253 to L262 for M4 as second point.
- **CV_3_**: the distance between M2 and M4 of the opposite subunits. To describe the CV we selected center of mass of all Cα atoms of residues from Y127 to P133 for M2 as first point and center of mass of all Cα atoms of residues from A253 to L262 for M4 as second point.
- **CV_4_**: the Ψ dihedral angle of Y175 residue located near S1 ion binding site.
- **CV_5_**: the Ψ dihedral angle of F284 residue located near S1 ion binding site.
- **CV_6_**: the presence of potassium ions at the S1 ion binding site. To describe the CV we used coordination numbers (continuous contacts) between potassium ions and center of S1 ion binding site.
- **CV_7_:** the presence of water molecules at the S1 ion binding site. To describe the CV we used coordination numbers (continuous contacts) between water molecules and center of S1 ion binding site.

To ensure thorough sampling and reliable convergence of the FES, we adopted the OneOPES replica-exchange setup comprising eight trajectories (Supplementary Table 1). Replica 0 acted as the convergence monitor, while the remaining seven replicas (1–7) were used to enhance exploration. All simulations employed OPES Explore on a defined set of collective variables, with the exploratory replicas progressively heated—up to 335 K—using the OPES Expanded (MultiT) scheme to facilitate sampling across hidden degrees of freedom^69^. Box dimensions and salt concentration for each system kept same (Supplementary Table 2).

All trajectories were analyzed with GROMACS tools together with PLUMED driver option^60^ and Python using MDAnalysis^70^ together with NumPy^71^ matplotlib^72^ and SciPy^73^. Distances, dihedral angles, coordination numbers and biases were calculated with PLUMED driver.

To calculate 1D and 2D free energy surfaces (FES), OPES biases were reweighted using weighted kernel density estimation. First 100 ns of each independent replica were skipped during FES calculations and considered as the equilibration of the system.

Molecular visualizations were rendered using PyMOL^52^. Figures including Fig. 3a, Fig. 4a and Fig. 6 were generated using Affinity Designer.

### Molecular biology and construct preparation

In this study the coding sequence of human TREK-2 (K_2P_10.1, *KCNK10*, GenBank accession number: NM_021161.5) was used. For TREK-2 channel expressed in *Xenopus* oocytes the coding sequence was subcloned into the oocyte expression vector pFAW. The A314P channel mutant was obtained by site-directed mutagenesis following the Quikchange method with custom oligonucleotides and verified by sequencing. Plasmid DNA was linearized with MluI and cRNA synthesized *in vitro* using the T7 HighYield^®^ T7 ARCA mRNA Synthesis Kit (Jena Bioscience, Germany).

### Animals and oocyte preparation

For this study, three female *Xenopus laevis* claw frogs were used to isolate oocytes. The investigation conforms to the guide for the Care and Use of laboratory Animals (NIH Publication 85-23) and were approved by the local ethics commission and permitted by the department for animal experimentation.

Ovarian lobes were obtained from frogs anesthetized with tricaine. Lobes were treated with type II collagenase (Sigma-Aldrich/Merck, Germany) in OR2 solution containing (in mM): 82.5 NaCl, 2 KCl, 1 MgCl_2_, 5 HEPES (pH 7.4 adjusted with (NaOH/HCl) for 2 h. Isolated oocytes were stored at 16.9° C in a solution containing (in mM): 54 NaCl, 30 KCl, 2.4 NaHCO_3_, 0.82 MgSO_4_ x 7 H_2_O, 0.41 CaCl_2_, 0.33 Ca(NO_3_)_2_ x 4 H_2_O and 7.5 TRIS (pH 7.4 adjusted with NaOH/HCl) and were manually defolliculated prior injection with cRNA.

### Two-electrode voltage-clamp (TEVC) recordings of Xenopus oocytes

Oocytes were injected with 2 ng of RNA for WT and mutant channels, respectively and incubated for 2 days at 16.9° C. Standard TEVC measurements were performed at room temperature (20 - 22° C) with a TEC-10CX TEVC system coupled to a HEKA EPC-9 patch clamp amplifier, and PatchMaster software (v2x92). Microelectrodes were fabricated from glass pipettes, back-filled with 3 M KCl, and had a resistance of 0.5 - 1.5 MΩ. Recordings were performed in ND96 buffer at pH 7.4 (96 mM NaCl, 2 mM KCl, 1 mM MgCl_2_, 1.8 mM CaCl_2_, 5 mM HEPES). Currents were recorded using 800 ms ramp protocols from -100 to +60 mV and a holding potential of -80 mV. All recorded ramps were analyzed at +40 mV.

### Inside-out patch-clamp recordings of Xenopus oocytes

Oocytes were injected with 2 ng of RNA for WT and mutant channels, respectively and incubated for 2 - 3 days at 16.9° C in a solution containing (mM): 54 NaCl, 30 KCl, 2.4 NaHCO_3_, 0.82 MgSO_4_ x 7 H_2_O, 0.41 CaCl_2_, 0.33 Ca(NO_3_)_2_ x 4 H_2_O and 7.5 TRIS (pH 7.4 adjusted with NaOH/HCl). Inside-out patch-clamp recordings under voltage-clamp conditions were performed at room temperature (20 - 22° C). Patch pipettes were made from thick-walled borosilicate glass GB 200TF-8P (Science Products, Germany), had resistances of 0.2 - 0.5 MΩ (tip diameter of 10 - 25 µm) and filled with a pipette solution (in mM): 4 KCl, 116 NMDG, 10 HEPES and 3.6 CaCl_2_ (pH 7.4 adjusted with KOH/HCl). Intracellular bath solutions and compounds were applied to the cytoplasmic side of excised patches via a gravity flow multi-barrel pipette system. Intracellular solution had the following composition (in mM): 120 KCl, 10 HEPES, 2 EGTA and 1 pyrophosphate (pH adjusted with KOH/HCl). Currents were recorded with an EPC10 amplifier (HEKA electronics, Germany) and sampled at 10 kHz or higher and filtered with 3 kHz (-3 dB) or higher as appropriate for sampling rate. Voltage ramp pulses (-80 mV to +80 mV) were applied from a holding potential of -80 mV for 800 ms with an inter-pulse interval of 9 s and were analyzed at a given voltage of +40 mV. The relative steady-state levels of inhibition for NFx were fitted with the following Hill equation:

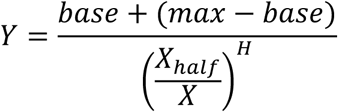

where *base* is the inhibited (zero) current, *max* is maximum current, *x* is the NFx concentration, *X_half_* is the value of concentration for half-maximal occupancy of NFx binding site and *H* is the Hill coefficient. Norfluoxetine (NFx) was purchased from Cayman Chemical (Biozol Diagnostics, Eching, Germany) and tetra-pentyl-ammonium (TPenA) from Sigma-Aldrich (Merck KGaA, Darmstadt, Germany). Compounds were stored as stock solution (50 - 100 mM) at -80° C and were diluted in intracellular bath solution to final concentrations prior to each measurement.

## Data availability

Data supporting the findings of this manuscript are available from the corresponding authors upon request. The trajectory data together with simulation parameters and initial structure of trajectories have been deposited in Zenodo under accession code https://doi.org/10.5281/zenodo.18622314.

## Code availability

Molecular dynamics simulation data were generated using the GROMACS software (2022.5)^59^ patched with PLUMED 2.8.5^60^. All trajectories were analyzed with GROMACS tools together with PLUMED driver option and Python using MDAnalysis together with NumPy matplotlib and SciPy.

## Acknowledgements

We greatly acknowledge Dr. Barth van Rossum for assistance with data visualization and Dr. Tillmann Utesch for technical support. This work was funded by the Leibniz-Forschungsinstitut für Molekulare Pharmakologie (FMP), the Deutsche Forschungsgemeinschaft (DFG, German Research Foundation) under Germanýs Excellence Strategy—EXC 2008/1 (UniSysCat) – 390540038 (to H.S.), and a Leibniz Collaborative Excellence project (grant number: K622/2024, to H.S., M.S., T.B.). The MD simulations were performed with resources provided by the Erlangen National High Performance Computing Center (NHR@FAU) and the high-performance computer “Lise” at the NHR Center (NHR@ZIB).

## Author Contributions Statement

H.S., V.R., M.S., and F.L.G. designed and directed the project with contributions from all authors. B.T. conducted the MD simulations, while the data was analyzed and interpreted by B.T., V.R., S.G., S.A., F.L.G, and H.S. M.S. and C.B. performed patch-clamp experiments and M.S., C.B., and T.B. analyzed the data. B.T. and M.S. prepared figures. B.T., H.S., V.R., M.S., and F.L.G wrote the manuscript with contributions from all authors.

## Figure Legends/Captions

**Supplementary Fig. 1.**
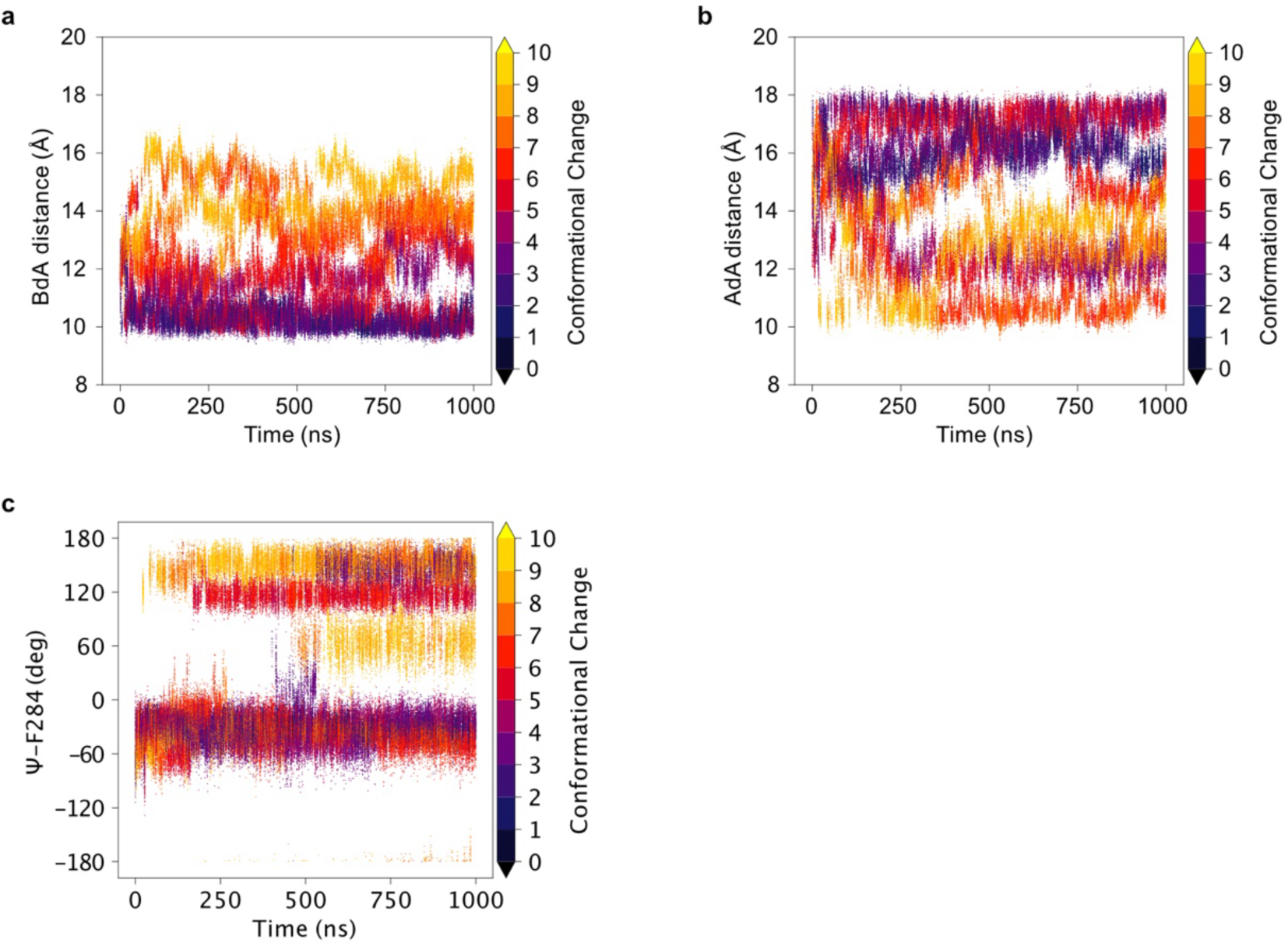
CV sampling in an apo TREK-2 OneOPES simulations. **(a,b)** Sampling of the M2-M4 distances as a function of time in replica 0, with data points colored according to the Path CV. **(c)** Sampling of the carbonyl movement of SF residue F284 as a function of time in replica 0, with data points colored according to the Path CV.

**Supplementary Fig. 2.**
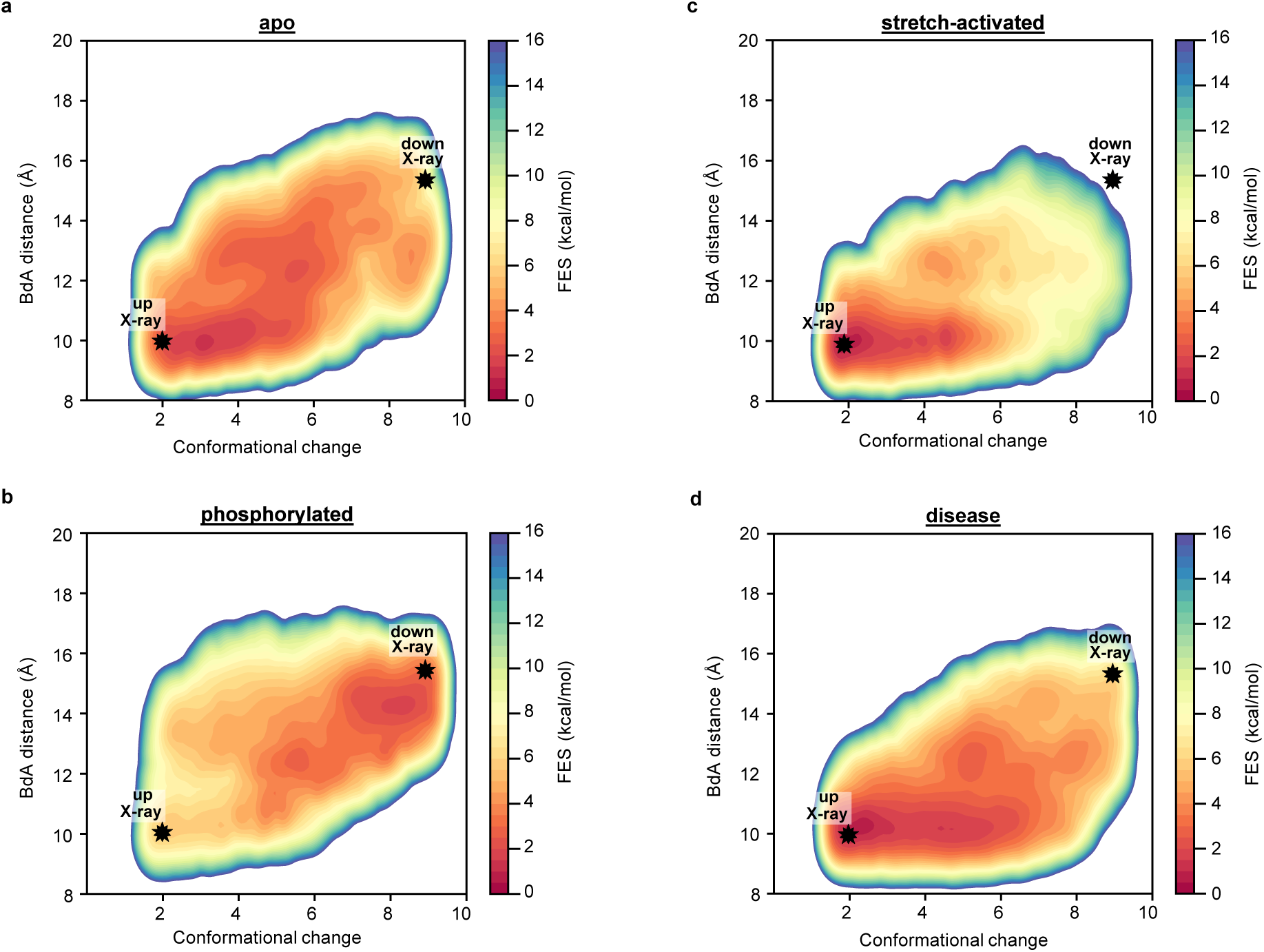
Analyses of the M4-pCt movement in TREK-2 OneOPES simulations. **(a)** Two-dimensional free energy surfaces projected along the Path CV and the auxiliary CV describing M4-M2 distance for apo, **(b)** phosphorylated, **(c)** membrane stretch and **(d)** disease simulations. The average of three independent replica for apo, phosphorylated and membrane stretch and five independent replicas for disease system were calculated for two-dimensional free energy surfaces. Black stars indicate the positions of the crystallographic TREK-2 up- and down-state structures mapped onto these coordinates, based on their measured path CV values and AdA distances.

**Supplementary Fig. 3.**
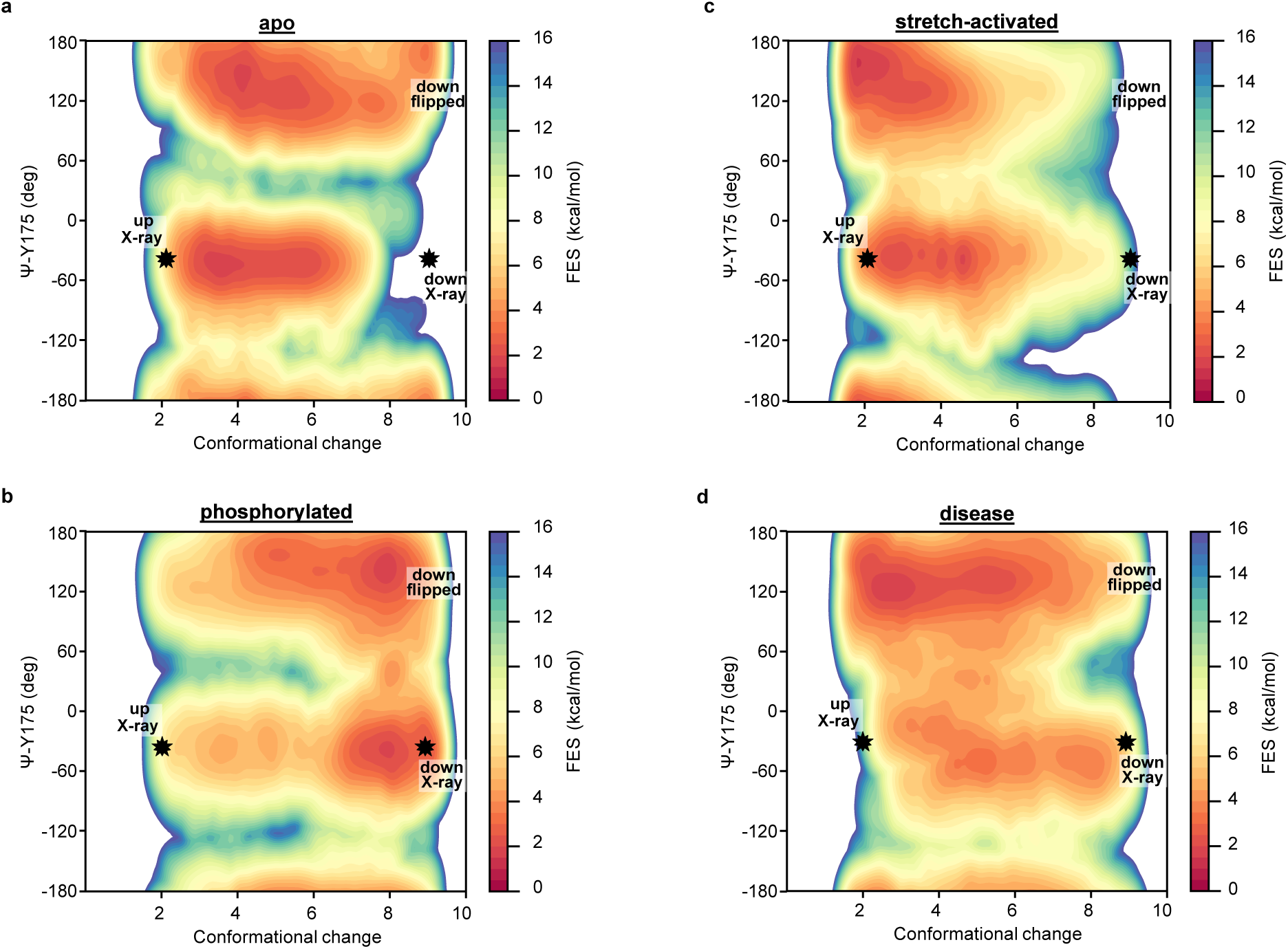
Analyses of the carbonyl conformation of Y175 in TREK-2 OneOPES simulations. **(a)** Two-dimensional free energy surfaces projected along the Path CV and the auxiliary CV describing carbonyl movement of Y175 SF residue for apo, **(b)** phosphorylated, **(c)** membrane stretch and **(d)** disease simulations. The average of three independent replica for apo, phosphorylated and membrane stretch and five independent replicas for disease system were calculated for two-dimensional free energy surfaces.

**Supplementary Fig. 4.**
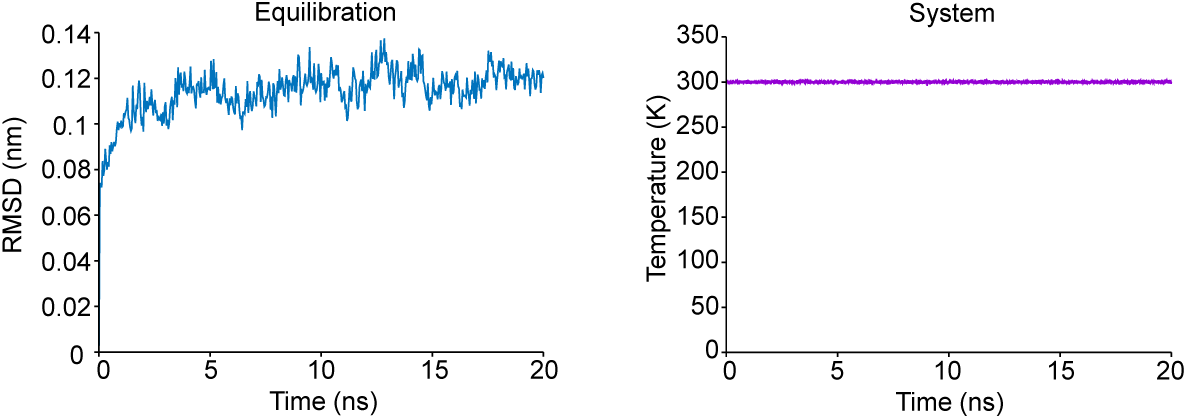
RMSD analysis of the MD data. The time course of root-mean-square deviation (RMSD) during the 20 ns equilibration and the temperature of the system during equilibration for structure of apo TREK-2.

**Supplementary Table 1.** CVs and parameters used in the TREK-2 OneOPES simulations. The rows labeled “OPES Explore” and “OPES MultiCV” indicate how the CVs were distributed across the different replicas. The “OPES MultiT” row reports the maximum temperature reached during the simulations, relative to the thermostat temperature (300 K).

| Replicas | 0 | 1 | 2 | 3 | 4 | 5 | 6 | 7 |
| --- | --- | --- | --- | --- | --- | --- | --- | --- |
| OPES Explore | p1.s | p1.s | p1.s | p1.s | p1.s | p1.s | p1.s | p1.s |
| OPES MultiCV<sub>1</sub> | – | m2bm4a | m2bm4a | m2bm4a | m2bm4a | m2bm4a | m2bm4a | m2bm4a |
| OPES MultiCV<sub>2</sub> | – | – | m4am2a | m4am2a | m4am2a | m4am2a | m4am2a | m4am2a |
| OPES MultiCV<sub>3</sub> | – | – | – | psiTa | psiTa | psiTa | psiTa | psiTa |
| OPES MultiCV<sub>4</sub> | – | – | – | – | psiFb | psiFb | psiFb | psiFb |
| OPES MultiCV<sub>5</sub> | – | – | – | – | – | V1, V2 | V1, V2 | V1, V2 |
| OPES MultiCV<sub>6</sub> | – | – | – | – | – | – | – | – |
| OPES MultiCV<sub>7</sub> | – | – | – | – | – | – | – | – |
| OPES MultiT | – | 301K | 303K | 306K | 310K | 317K | 325K | 335K |

**Supplementary Table 2.** Simulation details of each simulation setup.

| *System* | *Total N of atoms* | *Total N of H<sub>2</sub>O* | *Salt concentration* | *Total N of lipid composition (POPC 52 atoms)* | *Box dimensions (Å)* |
| --- | --- | --- | --- | --- | --- |
| *TREK-2 apo OneOPES* | 121627 | 32253 | 612 mM | 295 | 104.1x104.1x140.9 |
| *TREK-2 phosphorylated OneOPES* | 121629 | 32253 | 612 mM | 295 | 104.1x104.1x140.9 |
| *TREK-2 stretch OneOPES* | 121627 | 32253 | 612 mM | 295 | 104.1x104.1x140.9 |
| *TREK-2 disease mutant OneOPES* | 121635 | 32253 | 612 mM | 295 | 104.1x104.1x140.9 |

